# Uncovering the Genetic Blueprint of the UAE’s National Tree: Genomic Evidence to Guide *Prosopis cineraria* (L.) Druce Conservation

**DOI:** 10.1101/2025.07.15.663052

**Authors:** Anestis Gkanogiannis, Salama Rashed Obaid Almansoori, Maher Kabshawi, Mohammed Shahid, Saif Almansoori, Hifzur Rahman, Augusto Becerra Lopez-Lavalle

## Abstract

*Prosopis cineraria* (L.) Druce is a keystone tree species of the arid and semi-arid regions of the West and South Asia, with critical ecological, cultural, and consevation significance. In the United Arab Emirates (UAE) and other regions of the Arabian Peninsula this beneficial tree is called Ghaf. Despite its importance, genomic resources and population-level diversity data for the tree remain limited. Here, we present the first comprehensive population genomics study of Ghaf, based on whole-genome re-sequencing of 204 individual trees collected across the UAE.

Following Single Nucleotide Polymorphism (SNP) discovery and stringent filtering, we analyzed 57, 183 high-quality LD-pruned SNPs to assess population structure, diversity, and gene flow. Principal component analysis (PCA), sparse non-negative matrix factorization (sNMF), and discriminant analysis of principal components (DAPC) revealed four well-defined genetic clusters, broadly corresponding to geographic origins. Genetic diversity varied significantly among groups, with observed heterozygosity (Ho), inbreeding coefficients (F), and nucleotide diversity (π) showing strong population-specific trends.

Genome-wide Fixation index *F* ^ST^ scans identified multiple highly differentiated genomic regions, enriched for genes involved in stress response, transport, and signaling. Functional enrichment using Gene Ontology (GO), Kyoto Encyclopedia of Genes and Genomes (KEGG), and Pfam annotations indicated overrepresentation of protein kinase activity, ATP binding, and hormone signaling pathways. TreeMix analysis revealed gene flow into one of the genetic clusters from both others, suggesting historical admixture and geographic connectivity.

To support data interpretation and stakeholder engagement, we developed two web-based tools for interactive visualization of sample geolocation and genetic structure. This work provides foundational insights into the genetic landscape of *P. cineraria* (L.) Druce, supporting conservation planning and long-term monitoring in the region.

## 1 INTRODUCTION

The Ghaf tree (*Prosopis cineraria* (L.) Druce) is a symbol of endurance and cultural heritage across the arid landscapes of the Arabian Peninsula. In the United Arab Emirates (UAE), it holds national importance as the official national tree, recognized for its ecological functions in stabilizing sand dunes, enhancing soil fertility, and supporting native biodiversity. As a deep-rooted nitrogen-fixing legume, Ghaf plays a critical role in sustaining desert productivity under extreme heat and water scarcity.

Despite its ecological and cultural significance, Ghaf faces increasing threats from urbanization, habitat loss, and overgrazing. At the national level, *Prosopis cineraria* (L.) Druce is currently assessed as ‘Least Concern’ under the IUCN Red List criteria due to its relatively widespread distribution and persistence in protected areas (National Red List, 2019).

However, within the Emirate of Abu Dhabi, local assessments and conservation efforts by the Environment Agency–Abu Dhabi (EAD) indicate that the species is considered endangered due to localized pressures and habitat fragmentation (Environment Agency–Abu Dhabi, 2020).

Despite its ecological and cultural significance, *P. cineraria* (L.) Druce has been largely neglected in global genomic initiatives, and little is known about its population-level genetic variation. This knowledge gap is particularly pressing given the escalating pressures of climate change, habitat fragmentation, and land-use change across arid regions. Understanding the genetic diversity and structure of Ghaf is essential for evidence-based conservation, restoration, and management strategies (Frankham et al., 2002).

Here, we present the first population-scale genomic analysis of *P. cineraria* (L.) Druce, focusing on 204 individuals sampled from across the UAE. Our study is grounded in the conservation mission of the International Center for Biosaline Agriculture (ICBA), a regional leader in dryland agroecology and plant genetic resource management, and the Environment Agency–Abu Dhabi (EAD), the Middle East’s largest environmental regulator, which also plays a key role in plant genetic resource management. Using genome-wide SNPs obtained via Whole Genome Re-Sequencing (WGS), we aim to: (1) characterize the population structure of Ghaf and patterns of genetic diversity across its range in the UAE; (2) identify genomic regions under differentiation and potential local adaptation; and (3) investigate gene flow and historical connectivity among geographically distributed groups.

Our findings are framed not only by evolutionary and ecological questions, but also by national conservation goals, supporting seed sourcing programs, restoration ecology, and long-term genetic monitoring. Importantly, this research can help identify and prioritize high-value populations for ex-situ conservation efforts, including gene banking and seed banking, thereby safeguarding genetic diversity for future restoration and resilience initiatives. In addition, we introduce two custom-built, open-access web tools that allow conservation practitioners, government entities, and the public to interactively explore the genetic landscape of the Ghaf tree.

By generating the first genome-scale dataset for this iconic desert species, our work establishes a critical foundation for conservation genomics in the Arabian Peninsula and highlights the value of integrating population genomics, spatial ecology, and digital tools to inform dryland biodiversity stewardship.

## 2 MATERIALS AND METHODS

### 2.1 Sample Collection, SNP Calling and Filtering

A total of 211 *Prosopis cineraria* (L.) Druce (Ghaf tree) individuals were collected across multiple locations in the United Arab Emirates, covering major environmental and geographic zones.

The identification of Ghaf tree is carried out using a combination of comprehensive flora reference books, detailed monographic descriptions, and confirmation through comparison with herbarium specimens from the EAD herbarium collection. These resources allow for accurate identification in natural habitats by examining distinctive features, including foliage, bark texture, and overall morphology. Identified specimens are then deposited in the EAD herbarium for future reference and verification.

The collecting areas and routes to be followed were determined according to the geographical areas of the UAE, the coastal lowlands in the west and east, the Western dune plains, the central desert, the alluvial plain, and the mountain belt.

In this phase of collecting, eight expeditions were launched, covering most parts of the UAE. The sampling strategy depended on the population size and variation in habitat diversity. In general, for widely distributed species, stratified random sampling was adopted, and not less than 10 trees were sampled from each collecting site. For the ecotypes that were very rare and found in very small numbers, all the plants available at the site were sampled. In the absence of any morphological variation, samples of an accession collected from similar ecological habitats within each geographic area were pooled into one, irrespective of the distance between the collecting sites. After being collected into plastic containers, the leaf samples were transferred immediately to liquid nitrogen for preservation and later for DNA extraction.

Upon identification, the collection of Ghaf tree leaves follows a structured protocol to ensure sample integrity. Leaves are carefully plucked using sterilized gloves to avoid contamination, and each leaf sample is placed into 50 ml Falcon tubes, which are clearly labeled with a unique identifier to maintain traceability. The exact location of each tree is recorded using GPS devices and registered in the EAD Geographic Information System (GIS) database, ensuring traceability of the samples and contributing to ecological studies and conservation efforts.

Following collection, the preservation and preparation of the Ghaf tree samples for analysis involve several steps. Upon collection, the Falcon tubes containing the samples are immediately stored in a dry ice box to prevent the degradation of biological material and preserve the samples’ viability for subsequent analysis. The samples are then transported under controlled conditions to ICBA’s Desert Life Science Laboratory (DLSL), where the preservation process continues. At the laboratory, the samples undergo preparatory procedures, including washing, drying, and grinding, to extract the necessary compounds for analysis. These steps follow standardized protocols to ensure consistency and reliability in the results. This systematic methodology guarantees the integrity of the Ghaf tree samples throughout the process, enabling accurate and reliable analysis.

DNA was extracted from flash-frozen leaf samples using a modified CTAB method (Porebski et al., 1997). The purity and quality of the isolated DNA were verified by agarose gel electrophoresis on 0.8% agarose gels, and the concentration was determined on a Qubit 4 fluorometer using Qubit Broad Range Assay Kit (Thermo Fisher Scientific Inc., Waltham, MA, USA). One microgram of genomic DNA was mechanically fragmented to an average size of 250 bp using the Covaris® M220 Focused-ultrasonicator™ (Covaris, Woburn, MA, USA) and the size selection of fragmented DNA was done using MGIEasy DNA-Clean beads (MGI Tech, Shenzhen, China). A single-stranded circular DNA library was prepared using MGIEasy Universal DNA Library Prep Set Ver. 1.0 following the manufacturer’s standard protocol for a 250-bp insert size, followed by DNA nano ball (DNB) formation based on rolling circle amplification. The DNB was loaded into the flow cell (DNBSEQ-G400RS Sequencing Flow Cell Ver. 3.0) and cPAS-based 100-bp paired-end sequencing was performed with DNBSEQ-G400RS High-Throughput Sequencing Set Ver. 3.1 (MGI Tech).

Raw reads were demultiplexed and quality-checked using FastQC v0.11.9 (Andrews, 2010). Reads were trimmed and filtered using fastp v0.20.1 (Chen et al., 2018) and aligned to the *P. cineraria* (L.) Druce reference genome using BWA-MEM v0.7.17 (Li and Durbin, 2009). Duplicates were marked using Picard Tools v2.26.10 (Broad Institute, 2019). SNP calling was performed jointly across all samples using GATK HaplotypeCaller v4.5.0.0 (Poplin, 2017) in GVCF mode, followed by joint genotyping with GenotypeGVCFs. The resulting raw VCF were filtered with VCFtools v0.1.16 (Danecek, 2011) by selecting bi-allelic SNPs only with minimum site quality of QUAL ≥30; minimum genotype quality GQ ≥20; minimum depth per genotype DP ≥5; maximum missingness per site *<* 10%; and minor allele frequency MAF ≥0.05.

Linkage disequilibrium (LD) pruning was then applied using PLINK v1.90b6.22 (Chang, 2015) with a sliding window of 50 SNPs, a step size of 5, and an *r*^2^ threshold of 0.5. This yielded a final set of 57, 183 high-quality LD-pruned SNPs.

A subset of 114 non-clonal samples was defined based on clonal identity analysis with fastreeR v2.0.0 (Gkanogiannis and Bruls, 2016), and all downstream population structure and diversity analyses were performed both on the full dataset and the non-clonal subset, depending on the objective.

### 2.2 Population Structure Analyses

To investigate the genetic structure within the *Prosopis cineraria* (L.) Druce population, we employed sparse non-negative matrix factorization (sNMF) v1.2 (Frichot, 2014). Analyses were conducted using the LD-pruned SNP dataset of 57, 183 SNPs from 114 individuals.

We tested values of K (number of ancestral populations) ranging from *K* = 1 to *K* = 10, with 20 replicates per K. The optimal K was selected based on the cross-entropy criterion, which minimizes prediction error.

Admixture coefficients (Q-matrix) for each individual were visualized as stacked bar plots and integrated with geographic data using pie charts plotted on a map. To address sample overlaps at identical or near-identical GPS coordinates, we developed a flower expansion algorithm that dynamically separates overlapping pies, implemented in a custom interactive web app (International Center for Biosaline Agriculture, 2025).

In addition, individual assignments were compared to Discriminant Analysis of Principal Components (DAPC) that was conducted using the adegenet v2.1.11 (Jombart, 2008) package. Input genotypes were converted to a genlight object, and population clusters (*K* = 4) were defined based on sNMF assignment. DAPC was performed using 40 principal components and 3 discriminant functions. Posterior membership probabilities and discriminant loadings were extracted for comparison with sNMF results.

### 2.3 Genetic Diversity and Differentiation

To assess genetic diversity within and between groups, we calculated standard population genetic statistics using the LD-pruned, non-clonal dataset (114 individuals and 57, 183 SNPs).

Observed heterozygosity (Ho) and expected heterozygosity (He) were computed using PLINK v1.9 and custom R scripts. The inbreeding coefficient (F) was calculated as. A proxy for per-sample nucleotide diversity (π) was also estimated based on the observed heterozygosity, normalized by total loci.

To evaluate group-level differences in these diversity indices (Ho, He, F, π), we applied ANOVA followed by Tukey’s HSD for post-hoc comparisons. Visualizations included violin plots, overlaid scatterplots, and summary tables, all produced in R v4.3.3 (R Core Team, 2024) and Python v3.10.1 (Van Rossum and Drake, 2009) custom scripts.

Genes overlapping the top 1% *F* _ST_ windows were identified using the *P. cineraria* (L.) Druce reference GFF3 annotation (Sudalaimuthuasari, 2022) and submitted for functional enrichment analysis using eggNOG-mapper v2.1.12 (Cantalapiedra, 2021) on the v5.0.2 eggNOG database. Enrichment was conducted for Gene Ontology (GO) terms and KEGG pathways, with additional visualization using GOATOOLS v1.4.12 (Klopfenstein, 2018).

### 2.4 Functional Divergence and Annotation

To estimate genetic differentiation between groups, we calculated pairwise Weir and Cockerham’s Fixation index (*F*_ST_) using VCFtools v0.1.16, applied both genome-wide and in sliding windows of 50*kb* and 10*kb* step to scan for regions of elevated differentiation.

Windows falling in the top 1% of the genome-wide *F*_ST_ distribution were considered candidate regions of divergence. These windows were intersected with the *P. cineraria* (L.) Druce genome annotation in GFF3 format, to identify overlapping or nearby genes.

Genes overlapping high *F*_ST_ windows were extracted and annotated using eggNOG-mapper v2.1.12, referencing the eggNOG v5.0.2 database. Functional annotation included: Gene Ontology (Ashburner, 2000) terms (biological process, molecular function, cellular component); KEGG orthology (Kanehisa and Goto, 2000) and pathway assignments; and Pfam Mistry (2021), InterPro (Blum, 2021), and UniProt (The UniProt Consortium, 2021) functional domains.

GO term enrichment analysis was conducted using GOATOOLS with a background set of all annotated genes in the genome. Enriched GO categories were filtered at *FDR <* 0.05. KEGG pathways were summarized using KO-term abundance mapping to the ko00001.keg hierarchy, and top pathways were visualized with matplotlib v3.10.1 (Hunter, 2007).

### 2.5 Phylogenetic Analysis

To visualize genetic relationships among *Prosopis cineraria* (L.) Druce individuals, we constructed a neighbor-joining (NJ) phylogenetic tree based on pairwise genetic distances.

Distances were calculated from the LD-pruned SNP dataset with fastreeR v2.0.0 (Gkanogiannis and Bruls, 2016) that uses cosine genetic distance as the distance metric.

The resulting NJ tree was: exported in Newick format; visualized with the iTOL v6 (Letunic and Bork, 2024) online tool and a custom-built interactive web-based tree viewer (International Center for Biosaline Agriculture, 2025); and colored by sNMF group assignments for comparison with admixture results.

### 2.6 Multivariate and Clustering Analysis (DAPC)

To complement the sNMF population structure analysis, we performed Discriminant Analysis of Principal Components (DAPC) using the adegenet package in R.

We used the LD-pruned, non-clonal genotype dataset (57, 183 SNPs, 114 individuals) and followed this pipeline: SNP genotypes were imported and converted into a genlight object; group assignments were defined based on sNMF clustering at *K* = 4; Principal Component Analysis (PCA) was first applied for dimensionality reduction; the number of PCs (*n*.*pca* = 40) was chosen based on a-score optimization; and discriminant analysis retained 3 discriminant functions to separate the four clusters.

Posterior probabilities of group membership and SNP loadings were extracted for interpretation and cross-validation against sNMF results.

### 2.7 Gene Flow and Admixture

We explored gene flow between the inferred Ghaf groups using two complementary approaches:

TreeMix Analysis: we used TreeMix v1.13 (Pickrell and Pritchard, 2012) to infer population splits and migration events; the LD-pruned, non-clonal SNP dataset (57, 183 SNPs, 114 individuals) was converted to TreeMix input using PLINK and custom scripts; groups were defined using sNMF (*K* = 4), and allele frequency matrices were generated per group; we tested migration edges (m) from 0 to 5 and rooted the tree at either Group–2 or Group–4; tree topology, edge weights, and residual fit were visualized using custom Python and R scripts; and arrows indicate direction and strength of gene flow events.

ABBA–BABA: we used Dsuite v0.5r58 (Malinsky et al., 2020) to perform D–statistics (ABBA–BABA) tests on the unpruned, 204–sample VCF; a population assignment file mapped samples to groups and specified outgroup individuals; both global and per-chromosome tests were conducted, the statistics D, Z-score, p-value, *f*_4_-ratio, and site patterns (BBAA, ABBA, BABA) were computed; and per-chromosome results highlighted local introgression events.

### 2.8 Linkage Disequilibrium Decay

We evaluated linkage disequilibrium (LD) decay across the genome and among groups using PopLDdecay v3.43 (Zhang, 2019): LD was calculated on the LD–pruned, non–clonal dataset (114 samples; 57, 183 SNPs); analyses were performed per chromosome (PC1–PC15), per sNMF group (Group–1 to Group–4) or combined groups (ALL); and output files containing pairwise *r*^2^ values and distances were generated for each (group to chromosome) combination.

For visualization: a custom Python pipeline parsed the stat files, binned *r*^2^ by physical distance, and fit generalized additive models (GAMs) using the pygam v0.9.1 (Servén and Brummitt, 2018) library; LD decay curves were plotted for all groups together across all chromosomes and per chromosome (optionally selecting subsets of groups); and for each curve, we calculated the half-decay distance (bp) at which *r*^2^ dropped to 50% of its maximum.

## 3 RESULTS

### 3.1 SNP Filtering and Dataset Summary

Out of the 211 initially sequenced Ghaf samples, 204 passed quality filtering and were retained for downstream analyses. Seven samples with excessive missing data or low coverage were excluded.

Geographic coordinates were available for all retained samples, allowing integration with spatial analyses (Fig. 1). A subset of 114 samples was identified as non–clonal and used in downstream population genetic analyses requiring unrelated individuals.

**Figure 1.**
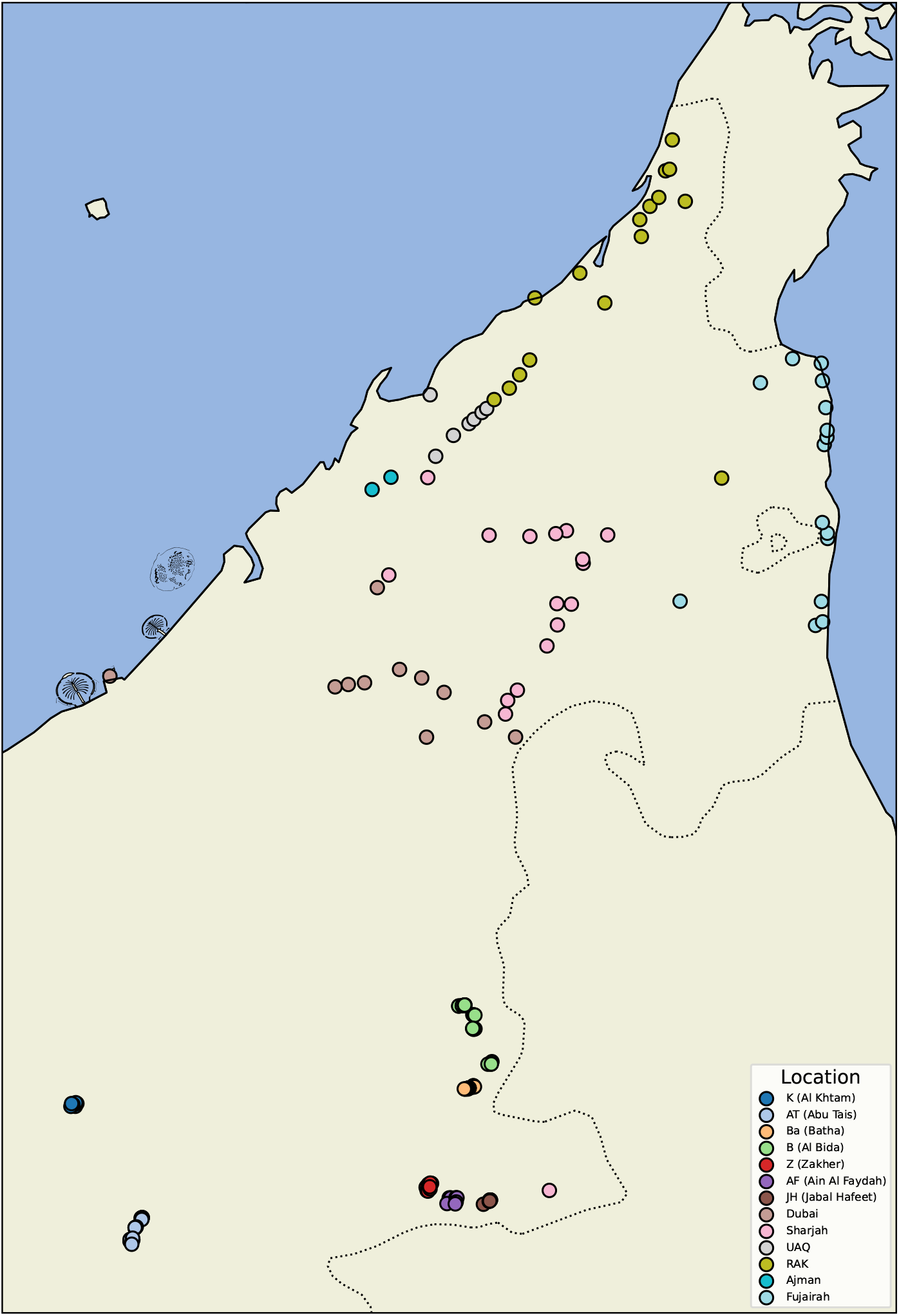
Sampling Map of *Prosopis cineraria* (L.) Druce Across the UAE. Geographic distribution of the 204 Ghaf tree samples used in this study. Points represent sampling locations across diverse ecological zones in the UAE, color-coded by site. The base map provides geographic context, highlighting widespread sampling coverage.

Whole-genome re-sequencing of 204 *Prosopis cineraria* (L.) Druce individuals yielded a total of 37, 302, 514 raw SNPs. After applying quality and filtering criteria to retain high-confidence bi-allelic variants, 432, 753 SNPs remained. Linkage disequilibrium (LD) pruning further reduced the dataset to 57, 183 SNPs, which formed the basis for all downstream population genomic analyses (Tab. 1).

**Table 1.**
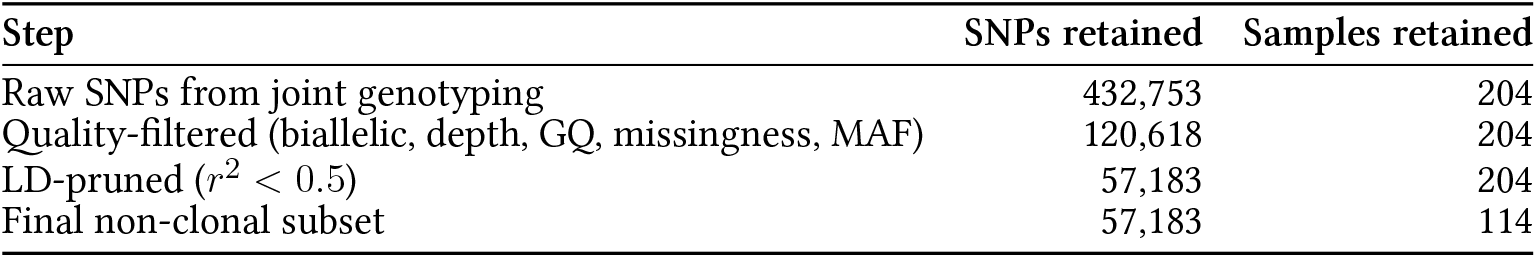
SNP filtering pipeline and resulting dataset sizes. Details of each filtering stage and final dataset sizes (SNPs and samples).

| Step | SNPs retained | Samples retained |
| --- | --- | --- |
| Raw SNPs from joint genotyping | 432,753 | 204 |
| Quality-filtered (biallelic, depth, GQ, missingness, MAF) | 120,618 | 204 |
| LD-pruned ($r^2 < 0.5$) | 57,183 | 204 |
| Final non-clonal subset | 57,183 | 114 |

### 3.2 Population Structure

Population structure analysis using sNMF revealed the presence of four major genetic clusters (*K* = 4). The optimal K was determined based on the minimum cross–entropy value, which sharply plateaued at *K* = 4 (Fig. 2A).

**Figure 2.**
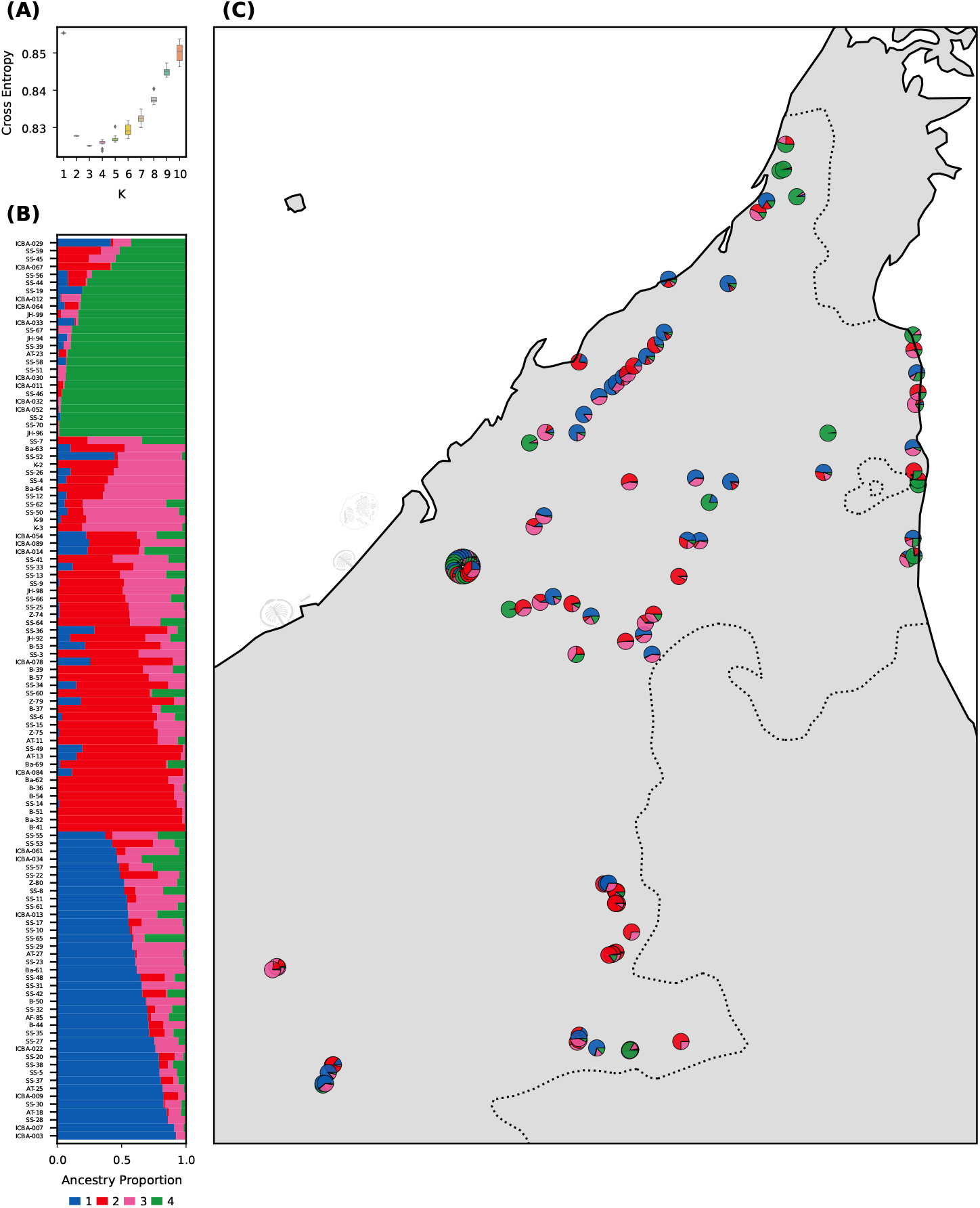
Population Structure Inference of *Prosopis cineraria* (L.) Druce via sNMF. Population structure inferred from 57, 183 LD-pruned SNPs using sNMF. Results indicate four ancestral clusters with distinct spatial patterns and admixture levels. (A) Cross-entropy criterion for *K* = 1 −10 shows optimal clustering at *K* = 4. (B) Q-matrix barplot showing individual ancestry proportions, sorted and grouped by major cluster. (C) Geographic map with pie charts of sNMF group membership per individual.

sNMF proportions showed variable levels of ancestral mixing across individuals, with some genetically homogeneous clusters and others showing significant admixture. Notably: Group–1 and Group–3 shared substantial ancestry; and Group–2 and Group–4 also showed affinity, with Group–4 displaying the most genetic differentiation.

sNMF patterns were visualized through stacked barplots (Fig. 2B), spatial pie charts, and geographic maps (Fig. 2C). A custom web-based visualization tool (International Center for Biosaline Agriculture, 2025) was developed to enable interactive exploration of the data, including geolocation and sNMF pie charts with overlapping samples resolved via flower-style expansions.

To further validate this structure, DAPC was applied to the same SNP dataset using sNMF–based groupings. The DAPC successfully discriminated the four clusters using just three discriminant functions (Fig. 3A). Posterior membership probabilities were high for most individuals (≥90%), with a small subset of samples showing admixed or ambiguous ancestry (Fig. 3B).

**Figure 3.**
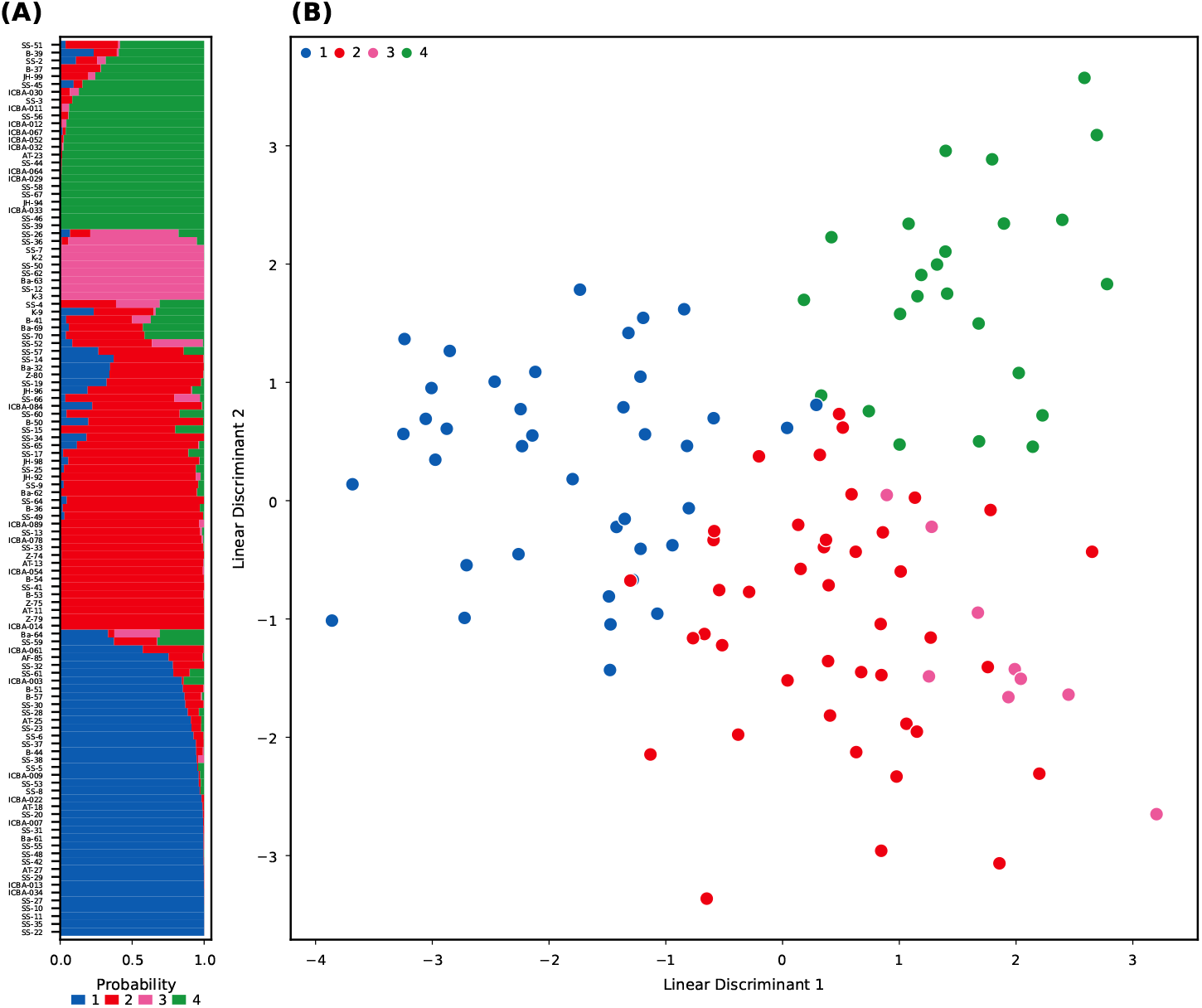
Discriminant Analysis of Principal Components (DAPC). DAPC confirms four genetically distinct clusters, with high membership probabilities. DAPC results closely mirror the sNMF clustering and show effective population separation. (A) Posterior membership probabilities of samples across groups. (B) DAPC scatter plot (LD1 vs LD2), with individuals colored by inferred group.

We also compared and confirmed individual assignments concordance between sNMF and DAPC using an alluvial plot (Supplementary Figure S1).

### 3.3 Genetic Diversity and Differentiation

We observed substantial differences in genetic diversity among the four sNMF-inferred groups (Fig. 4; Supplementary Figure S2): Group–4 showed significantly lower heterozygosity (Ho, He) and nucleotide diversity (π), along with higher inbreeding coefficient (F); and Group–1 and Group–3 displayed the highest diversity metrics, while Group–2 was intermediate.

**Figure 4.**
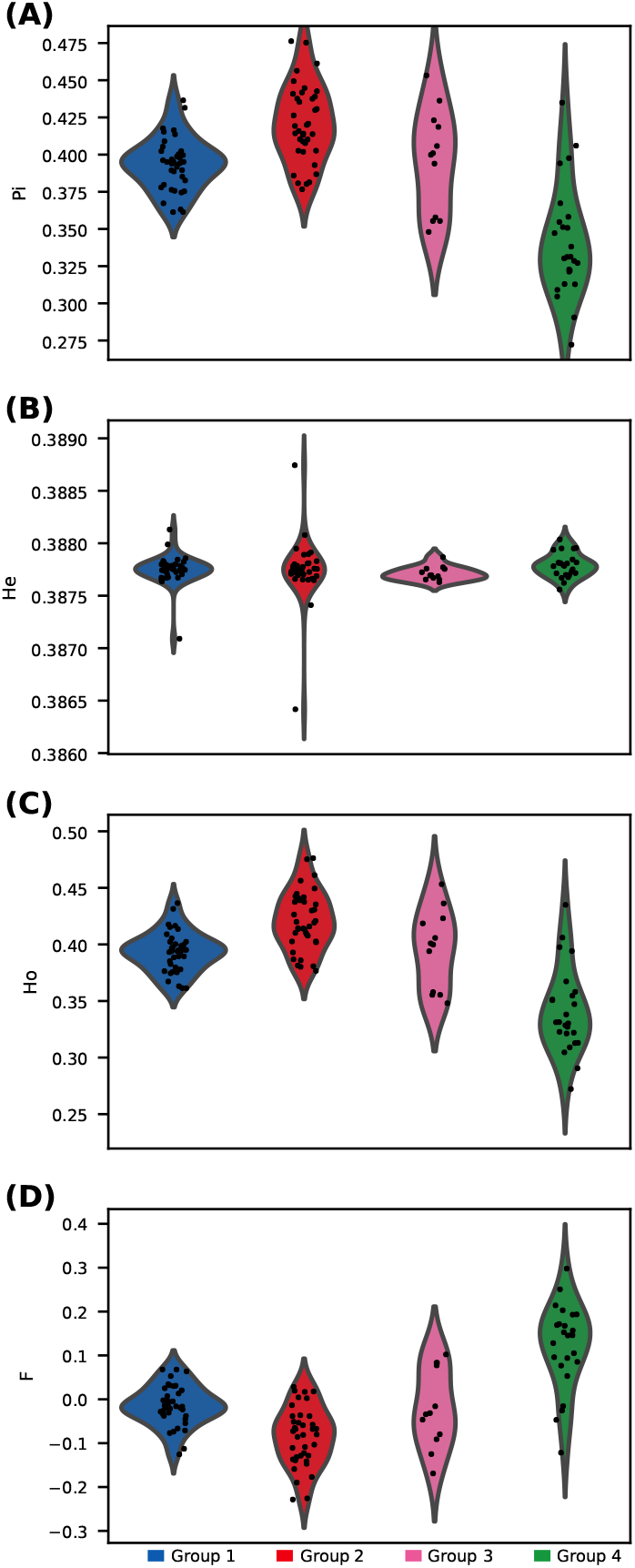
Genetic diversity across Ghaf groups. Violin plots with scatter overlays for four genetic diversity indices across sNMF groups. Group–4 shows markedly lower diversity and higher inbreeding. (A) Nucleotide diversity (π). (B) Expected heterozygosity (He). (C) Observed heterozygosity (Ho). (D) Inbreeding coefficient (F).

ANOVA tests followed by Tukey HSD confirmed that Group–4 differed significantly from the other groups for most diversity metrics.

Pairwise *F*_ST_ comparisons indicated moderate to strong differentiation, particularly involving Group– 4 (Tab. 2; Fig. 5):

**Table 2.**
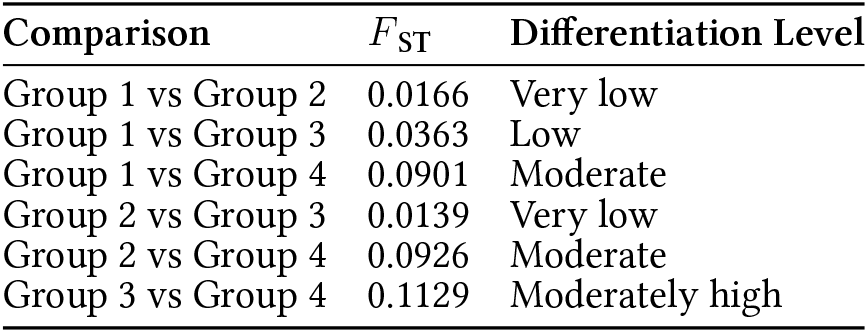
Pairwise *F*_ST_ matrix. Numeric *F*_ST_ values between all group combinations.

**Figure 5.**
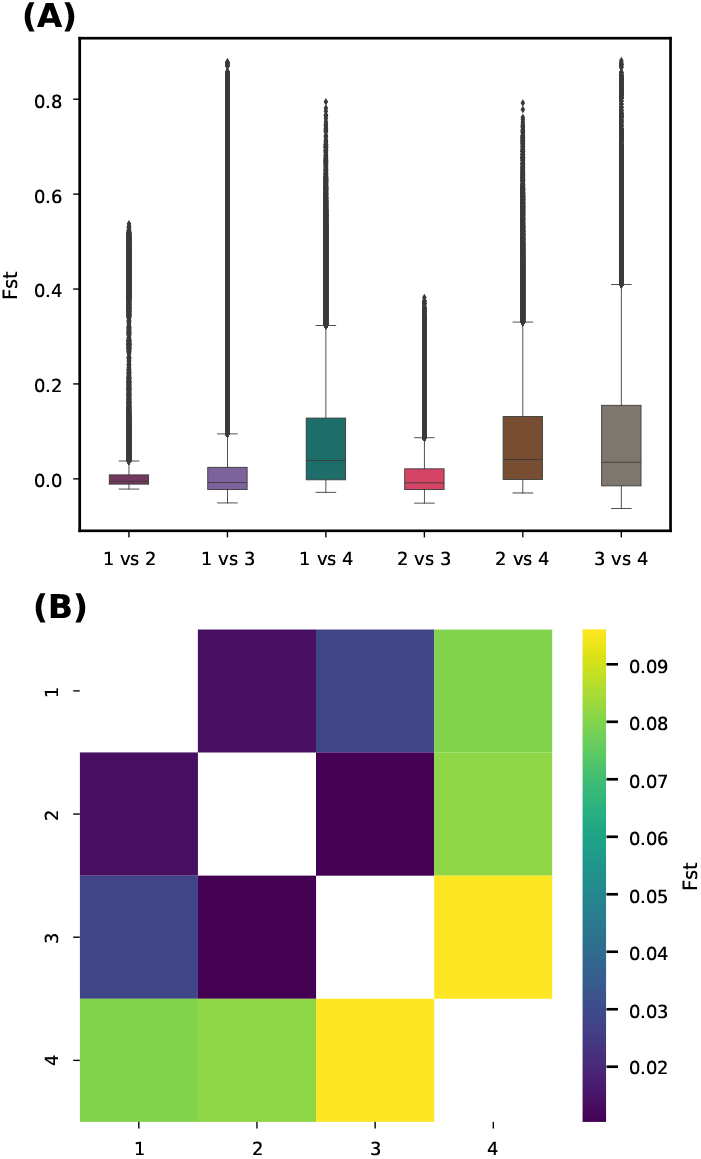
Summary of Pairwise *F*_ST_ Between Groups. Genetic differentiation is strongest between Group–4 and other groups, particularly Group-3 (*F*_ST_ = 0.113). Group–1 and Group–2 show low differentiation. (A) Boxplot distributions of SNP-wise *F*_ST_ for all six group comparisons. (B) Symmetric heatmap of mean pairwise *F*_ST_ values.

We also performed a genome-wide *F*_ST_ scan in sliding windows (50 kb, step 10 kb) to identify genomic regions under potential differentiation (Fig. 6; Supplementary Figure S3).

**Figure 6.**
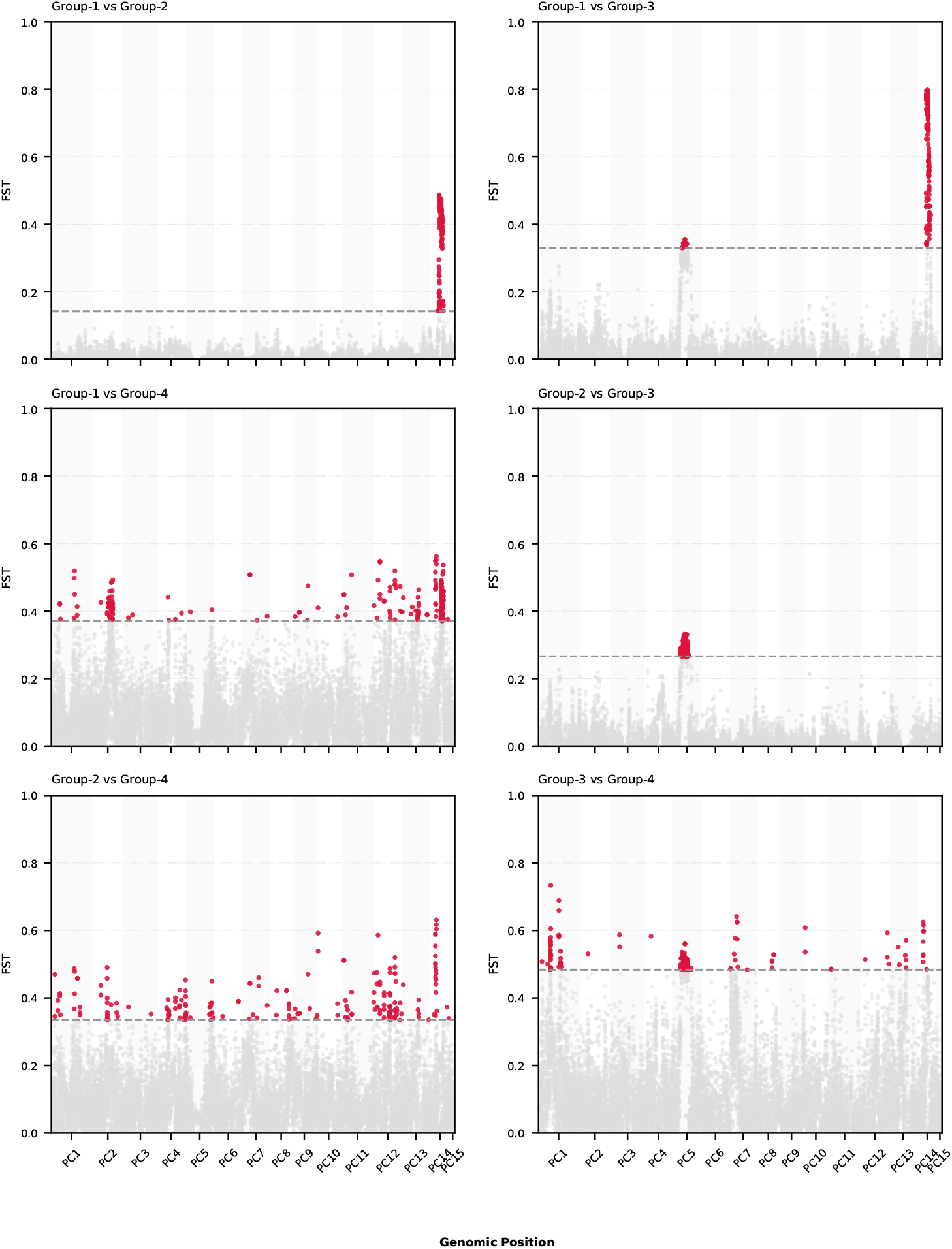
Genome-wide differentiation among Ghaf groups. Grid of Manhattan-style plots of windowed *F*ST (50 kb windows, 10 kb step) for all 6 pairwise group comparisons. The top 1% sliding window regions are highlighted Figure 7. Functional Enrichment of *Gkanogiannis et al*. Population Genomics of Ghaf Tree in UAE

**Figure 7.**
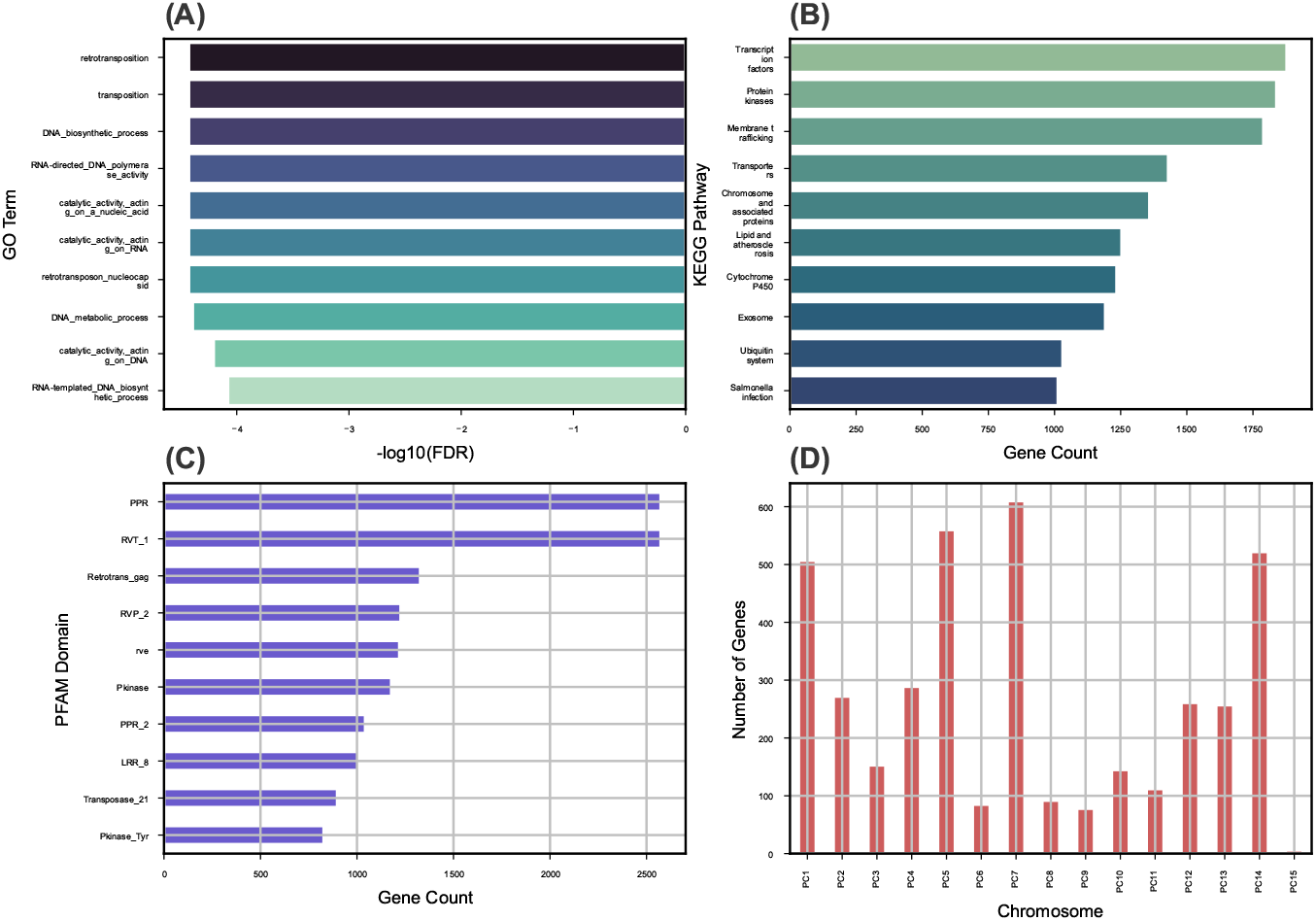
Functional Enrichment of High-*F*_ST_ Genes. High *F*_ST_ genes show enrichment for functions related to stress response, signaling, and metabolism. Divergent regions overlap with kinase domains, transporters, and retrotransposon activity. **(A)** Top enriched GO terms. **(B)** KEGG pathway enrichment. **(C)** PFAM domain abundance. (D) Top divergent genes by *F*_ST_ per chromosome.

These results confirm that Group–4 is genetically distinct, with lower intra-group diversity and higher divergence from the rest.

### 3.4 Functional Divergence and Enrichment Analysis

To link regions of high genetic differentiation to potential functional relevance, we extracted the top 1% *F*_ST_ windows (*±*2500 bp) and intersected them with gene models (Fig. 6).

A total of 3, 960 genes were found to contain or overlap high *F*_ST_ SNPs (Tab. 3). These genes were annotated using eggNOG-mapper (GO, KEGG, PFAM). Enrichment analysis (*FDR <* 0.05) showed functional enrichment in processes related to stress response, kinase activity, membrane localization and secondary metabolism.

Among these genes ∼85% received functional annotation through eggNOG-mapper, 780 genes were assigned one or more GO terms, and 326 genes mapped to KEGG orthologs (KOs), representing 118 distinct KEGG pathways.

Gene Ontology (GO) enrichment analysis revealed several overrepresented biological processes, including response to stress, signal transduction, and protein phosphorylation. At the molecular function level, enriched terms included kinase activity, ATP binding, and metal ion binding.

KEGG pathway analysis identified several significantly enriched and high-abundance pathways, notably plant hormone signal transduction, MAPK signaling, biosynthesis of secondary metabolites, and pathways associated with environmental adaptation.

These findings suggest that genes under high differentiation among Ghaf genetic clusters are involved in stress response, signaling, and local adaptation processes, consistent with ecological divergence in the UAE desert environment.

### 3.5 Phylogenetic Analysis

The Neighbor-Joining (NJ) tree (Fig. 8) constructed from 57, 183 LD–pruned SNPs, revealed clear clustering of individuals consistent with the four sNMF groups. Group–1 and Group–3 formed a cohesive clade, inticating close genetic proximity. In contrast, Group–2 and Group–4 clustered separately, with Group–4 forming a more distinct and more divergent branch, consistent with its elevated *F*_ST_ and higher inbreeding coefficients.

**Figure 8.**
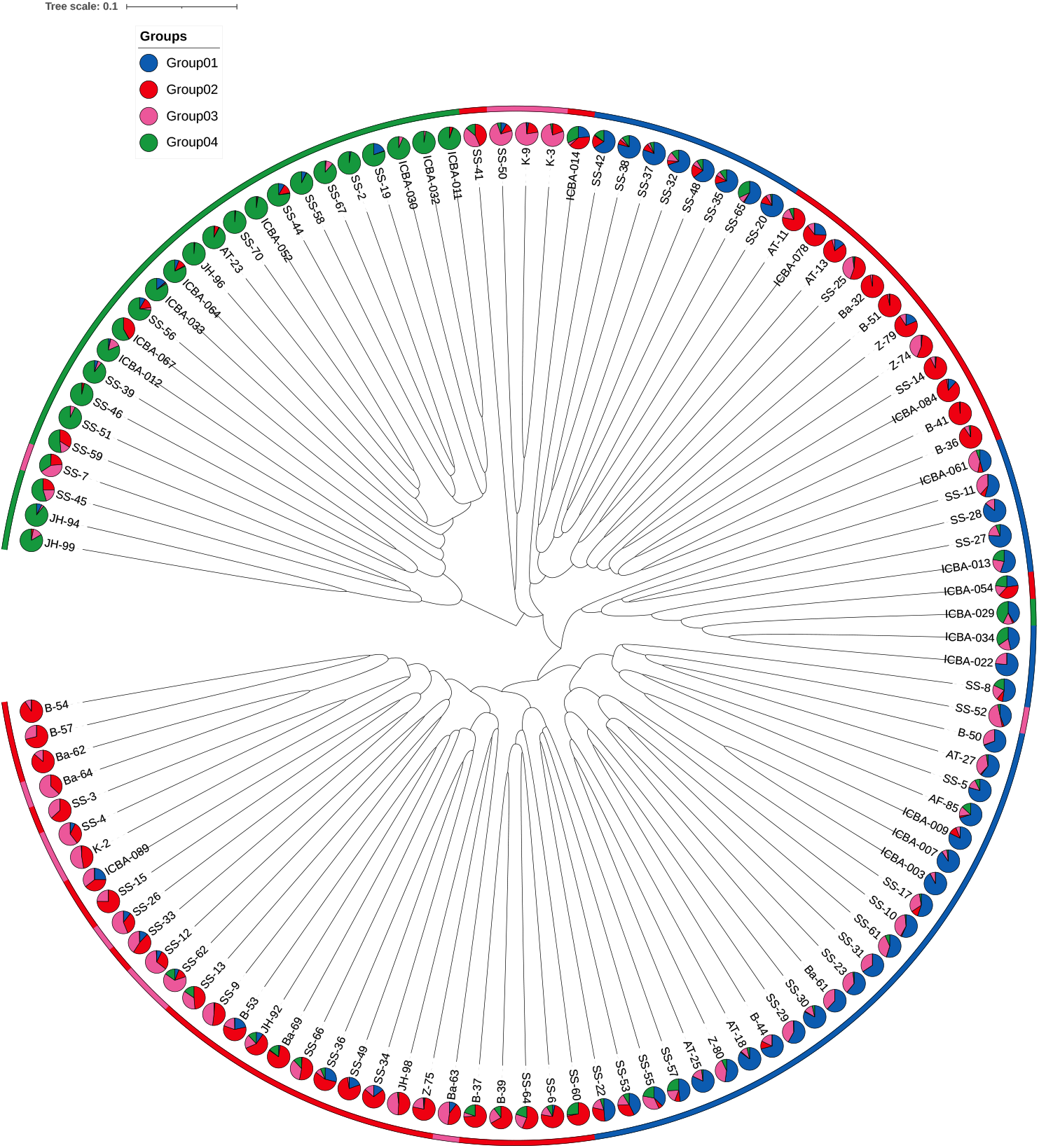
Phylogenetic relationships among Ghaf individuals. Neighbor-joining tree colored by sNMF group assignment. Branch structure confirms population subdivisions and supports sNMF/DAPC clustering.

The phylogenetic structure corroborates both the sNMF and DAPC clustering results, further supporting the presence of four distinct genetic lineages within the UAE *P. cineraria* (L.) Druce population.

An interactive web-based tree viewer (International Center for Biosaline Agriculture, 2025) allows dynamic exploration of the tree and access to individual sample metadata.

### 3.6 Gene Flow and Admixture

#### 3.6.1 TreeMix Results

Population splits and migration edges were inferred using TreeMix across the four sNMF–defined distinct genetic lineages within the UAE *P. cineraria* (L.) Druce population (Supplementary Figure S4). All tested models converged on similar topologies: Group–1 and Group–3 clustered together, while Group–2 and Group–4 formed a separate branch. Notably, a single migration edge consistently emerged between Group–1 and Group–2, suggesting historical gene flow between them.

#### 3.6.2 ABBA-BABA (Dsuite)

Genome-wide analysis using the ABBA–BABA (D-statistic) test revealed no significant global signal of introgression, indicating limited overall gene flow. However, chromosome–level tests uncovered localized signatures of gene flow, particularly on chromosomes PC1, PC5, PC8, PC9, PC12, and PC14. In several of these chromosomes, Z–scores exceeded 2, supporting hypotheses of introgression at specific loci (Supplementary Figure S5). These results provide independent evidence for localized gene flow, despite the overall well–differentiated population structure.

### 3.7 Linkage Disequilibrium Decay

LD decay patterns varied among groups and across chromosomes. Group–4 consistently exhibited steeper decay curves and shorter half–decay distances, likely reflecting a smaller effective population size or a more recent bottleneck (Fig. 9). In contrast, Group–1 and Group–3 displayed shallower LD decay patterns, consistent with larger or more stable demographic histories. Chromosome–level LD decay plots further highlighted genomic heterogeneity in LD patterns, with chromosomes such as PC1, PC5, PC8, and PC14 showing evidence of longer–range LD (Supplementary Figure S6).

**Figure 9.**
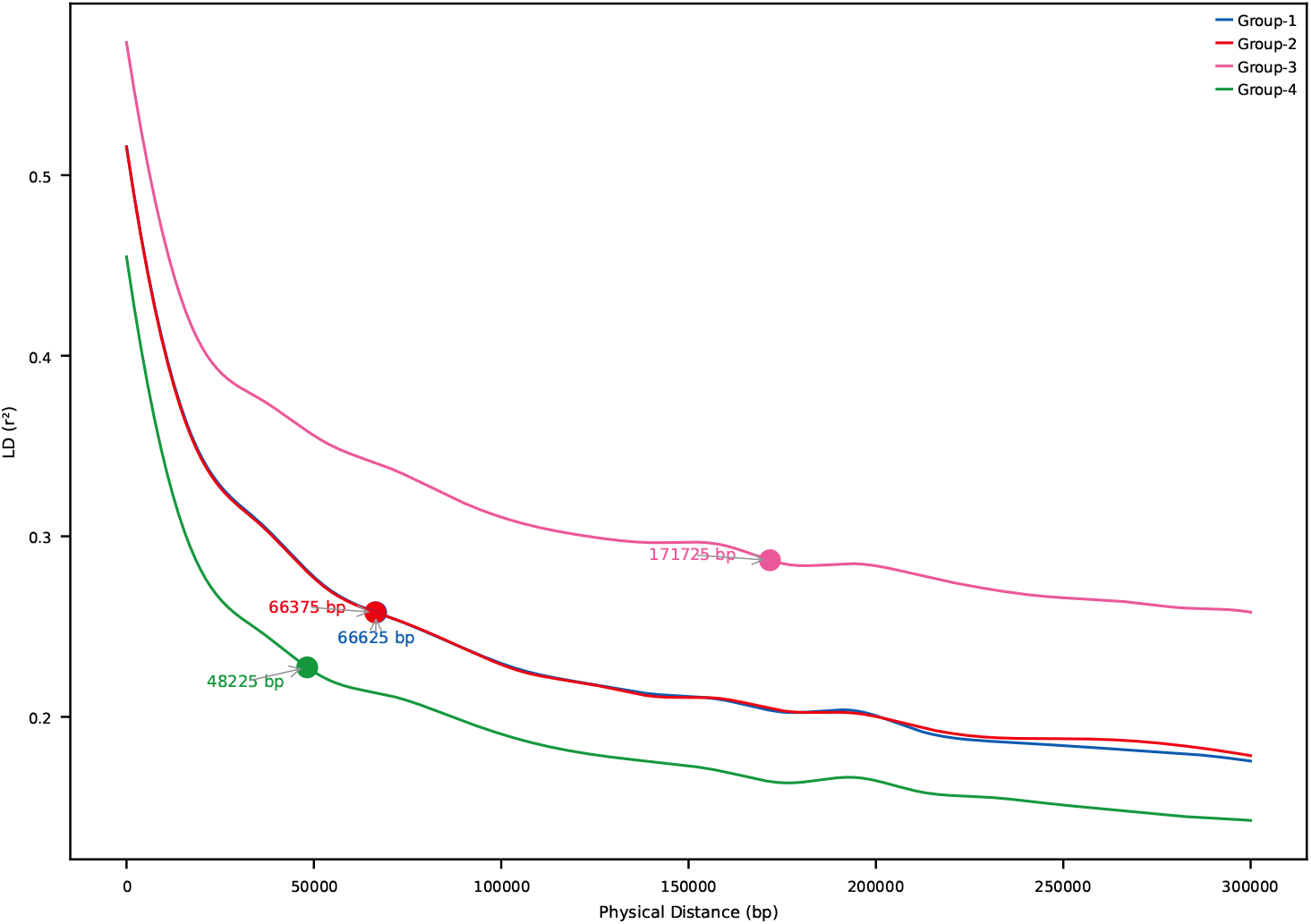
LD decay by group. Group–4 shows steeper LD decay, whereas Group–3 showed a shallower decay, suggesting a more stable population history.

## 4 DISCUSSION

Our study presents the first population-scale genomic assessment of *Prosopis cineraria* (L.) Druce (Ghaf tree) in the United Arab Emirates, leveraging genome–wide SNPs, spatial coordinates, and functional annotations. The findings illuminate how geography, genetic drift, and localized selection have shaped the species’ genomic landscape across its arid habitat.

### 4.1 Four Genetic Clusters Reflect Geography and Admixture

We identified four well–supported genetic clusters using sNMF and DAPC (Fig. 2; Fig. 3), consistent with subtle isolation–by–distance and regionally distinct demes. Groups 1 and 3, and Groups 2 and 4, formed two broader genetic meta–clusters with partial admixture, likely reflecting historical connectivity and subsequent drift.

Groups 1, 2, and 3 are closely related, with *F*_ST_ *<* 0.04 between any pair, consistent with low–to– moderate structure and potential gene flow or recent divergence. Group–4 is clearly distinct; it shows moderate–to–high differentiation from all others (*F*_ST_ *>* 0.09). The highest value of *F*_ST_0.1129 against Group–3, indicating relatively strong divergence. The progression of *F*_ST_ values suggests that Group– 4 may represent a geographically or ecologically isolated lineage, possibly under distinct selective pressures or with reduced gene flow.

These patterns align with biogeographic expectations for a long–lived, insect–pollinated desert tree in fragmented habitats (Gairola et al., 2019). The availability of GPS-linked samples allowed spatial interpretation of genetic clusters (Fig. 2C), with localized structure and pockets of genetic homogeneity near protected or isolated sites.

### 4.2 Group–4: A Genetically Distinct and Vulnerable Lineage

Group–4 emerged as a genetically depauperate and differentiated cluster, showing lower heterozygosity, higher inbreeding coefficients (Fig. 4), and strong pairwise *F*_ST_ values (Fig. 5). This suggests isolation, drift, and potential bottlenecks, a warning signal for long-term viability if unmonitored (Frankham et al., 2002).

It contains samples like SS–44, SS–45, SS–67, etc. that are located far northeast, in the Hajar Mountains, the coastal Ras Al Khaimah, and northern Fujairah region. Respectively, many ICBA–XXX samples in this group are also genetically isolated despite being collected at the same ICBA site, suggesting a unique lineage possibly due to clonal propagation or distinct origin. This group may represent either an ancestral lineage preserved in the northeast, or a human–mediated introduction. Its strong divergence suggests limited gene flow with the others.

Such patterns are not uncommon in desert trees with restricted seed dispersal and long generation times (Thomas et al., 2010). Based on our survey and field assessments (data not presented), we speculate that Ghaf (*Prosopis cineraria* (L.) Druce) populations in mountainous and near-mountain areas exhibit subtle morphological differences compared to those in sand sheet and sand dune environments. These differences include variations in leaf structure, trunk growth, and other traits, which may reflect habitat– specific adaptations. Given its uniqueness, Group–4 could represent an Evolutionarily Significant Unit (ESU) and should be prioritized in UAE conservation strategies

### 4.3 Signals of Local Adaptation in Differentiated Genomic Regions

Sliding–window *F*_ST_ scans (Fig. 6) revealed several highly differentiated regions between group pairs, particularly between Group–1 and Group–4. Enrichment analyses of genes overlapping these high– *F*_ST_ windows uncovered biological functions potentially linked to desert adaptation, including stress response, kinase signaling, and secondary metabolite biosynthesis (Fig. 7; Tab. 3).

**Table 3.** High-*F*_ST_ regions per group comparison, collectively overlapping 3, 960 genes. The number of SNPs found in the top 1% *F*_ST_ regions per group comparison. Collapsed number of genes found spanning these SNPs (*±*2500 bp).

| Comparison | Outlier SNP Count | Threshold $F_{ST}$ | Outlier Gene Count |
| --- | --- | --- | --- |
| Group 1 vs Group 2 | 4349 | 0.4756 | 235 |
| Group 1 vs Group 3 | 4566 | 0.7959 | 220 |
| Group 1 vs Group 4 | 4290 | 0.4780 | 1283 |
| Group 2 vs Group 3 | 4286 | 0.2728 | 422 |
| Group 2 vs Group 4 | 4290 | 0.4345 | 1796 |
| Group 3 vs Group 4 | 4276 | 0.5690 | 1844 |

GO enrichment results strongly suggest that divergent genomic regions between Ghaf subpopulations are enriched in transposable element-related functions, especially retrotransposons and genes involved in reverse transcriptase activity, polymerases and nucleic acid remodeling. This enrichment in retrotransposon activity may signal active genome reshaping in Ghaf tree populations. This could be linked to adaptation, stress responses, or epigenetic regulation, especially relevant in the harsh arid environments of the UAE. This further might be indicating a mechanism for generating genetic diversity in otherwise clonally propagated populations.

These findings echo in previous ecological and transcriptomic studies suggesting *P. cineraria* (L.) Druce has evolved specific physiological responses to water scarcity, heat, and salinity. The identification (Sehgal et al., 2025) of candidate genes and pathways expands our understanding of the molecular basis of resilience in desert trees.

### 4.4 Evidence of Asymmetric Gene Flow

TreeMix analysis (Supplementary Figure S4) consistently inferred a migration edge from Group–1 to Group–2, a signal that was mirrored in ABBA-BABA tests showing elevated D–statistics in several chromosomes (Supplementary Figure S5). While global gene flow signals were weak, chromosome-specific results suggest historical episodes of introgression, possibly linked to shared environmental pressures or secondary contact zones.

Such localized introgression may reflect past hybrid zones or stepping–stone gene flow facilitated by intermittent connectivity during wetter climatic periods (Zachos, 2001). *Prosopis cineraria* may represent a remnant species from a significantly wetter climatic period in the distant past. Its continued presence in the region underscores its resilience in the face of progressive aridification over the past approximately 5, 000 years (Brown and Feulner, 2024); (Paparella and Burt, 2024).

### 4.5 Contrasting LD Decay Profiles Reflect Population History

LD decay analysis revealed notable differences across groups (Fig. 9; Supplementary Figure S6). Group– 4 again showed the most rapid decay, suggesting more recombination events and smaller LD blocks, consistent with a longer or more stable demographic history. In contrast, groups with broader or more admixed ancestries (e.g., Group–1 and Group–3) retained longer LD blocks, which may be indicative of past expansions or reduced effective recombination due to structure.

These differences in LD patterns should inform future association mapping efforts and provide insights into the evolutionary forces shaping genome-wide linkage in arid–adapted trees.

### 4.6 Implications for Conservation

As the national tree of the United Arab Emirates, *Prosopis cineraria* (L.) Druce holds ecological, cultural, and symbolic significance. It is recognized as a keystone species for its role in dune stabilization, shade provisioning, and nitrogen cycling, aligning with the sustainability goals outlined in UAE Vision 2031. Our findings have direct implications for enhancing ecosystem resilience amid climate change. They support targeted conservation and restoration strategies that contribute to the development of climate– resilient landscapes across the UAE.

First, the four genetically distinct clusters identified through our analyses should be considered separate conservation units. Notably, Group–4 appears particularly divergent and may require targeted in situ protection to avoid genetic swamping and preserve its unique lineage.

Second, the candidate genes and pathways identified as being under selection (Tab. 3 and Supplementary Data) highlight sources of adaptive genetic variation. These loci could be leveraged to guide assisted gene flow or targeted restoration, particularly in regions experiencing increasing desertification or salinity stress.

Third, our results can inform seed sourcing strategies for diversity maintenance and support regional replanting efforts, including national programs such as the Ghaf National Initiative and legacy reforestation efforts following Expo 2020.

Finally, the development of interactive genomic maps and phylogenetic tree visualizers ((International Center for Biosaline Agriculture, 2025); Fig. 2C; Fig. 8) provides practical tools for long-term monitoring and decision–making. These tools could support scalable implementation by UAE environmental authorities such as the Environment Agency Abu Dhabi (EAD).

Altogether, our study illustrates how population genomic data can align with national policy objectives and contribute to evidence-based biodiversity stewardship in arid and semi–arid regions, particularly in biodiversity hotspots like the Arabian Peninsula, where conservation genomics remains an emerging discipline.

### 4.7 Future Directions

This study lays a strong foundation for continued genomic, ecological, and conservation research on *Prosopis cineraria* (L.) Druce. Several avenues merit further exploration. High–coverage whole–genome resequencing of representative individuals would enable the detection of regulatory and non–coding variants potentially involved in local adaptation. Functional validation of candidate adaptive genes could be achieved through transcriptomic profiling under stress conditions. Additionally, defining seed zones and developing assisted gene flow strategies will be essential, particularly for genetically depauperate populations such as Group–4. Comparative genomic analyses with other *Prosopis* species, or even unrelated desert–adapted trees from the region could shed light on convergent evolutionary responses to arid environments.

Expanding the current framework to include *P. cineraria* (L.) Druce populations from neighboring countries (e.g., Oman) would provide a broader biogeographic perspective. Finally, integrating genomic insights into citizen science initiatives and national planting campaigns would help promote long–term resilience in reforested or restored habitats.

## 5 CONCLUSION

This study provides a comprehensive genomic snapshot of *P. cineraria* (L.) Druce populations across the United Arab Emirates. Our analyses revealed clear population structure influenced by geographic separation and admixture, alongside the presence of a genetically distinct and less diverse cluster (Group–4), which may warrant targeted conservation efforts. We identified genomic regions with strong differentiation, many of which are enriched for functions related to environmental adaptation. Patterns of historical gene flow were detected at the chromosome level, and variation in linkage disequilibrium decay among groups reflected differences in demographic history and recombination dynamics.

Beyond its scientific contribution, this work offers critical insights for the evidence–based conservation of *P. cineraria* (L.) Druce, the national tree of the UAE. It also expands the genomic toolkit available for studying desert–adapted trees, thereby supporting regional strategies for biodiversity management and climate resilience in arid ecosystems.

## DATA AVAILABILITY STATEMENT

All raw sequencing data have been deposited in INSDC repository under accession number PRJEB82449. Raw VCFs, sample metadata and other project related files have been deposited in Zenodo repository 10.5281/zenodo.15716843.

## AUTHOR CONTRIBUTIONS

AG: Methodology, Software, Formal analysis, Data curation, Writing–original draft, Writing–review & editing, Visualization. SROA: Conceptualization, Investigation, Writing–review & editing. MK: Writing–review & editing, Supervision. MS: Investigation, Data curation, Writing–review & editing. SA: Investigation, Data curation, Writing–review & editing. HR: Conceptualization, Investigation, Data curation, Writing–review & editing. ABLL: Conceptualization, Writing–review & editing, Supervision.

## FUNDING

Funding was received from ICBA’s core donor, the UAE Government, under Project code ICBA.079.

ACKNOWLEDGMENTS

We thank the field teams from ICBA and EAD for logistical support and assistance with sample collection.

## ETHICS AND PERMITS

All sample collection adhered to relevant national biodiversity and conservation guidelines. In addition, sample collection within the Emirate of Abu Dhabi was conducted under the authority of the Environment Agency - Abu Dhabi.

