## Supplementary Material for "Uncovering the Genetic Blueprint of the UAE’s National Tree: Genomic Evidence to Guide *Prosopis cineraria* (L.) Druce Conservation"

### 1 SUPPLEMENTARY FIGURES

The following supplementary figures provide additional visual and analytical insights supporting the main findings of this study. All supplementary content is available in the online repository associated with this manuscript.

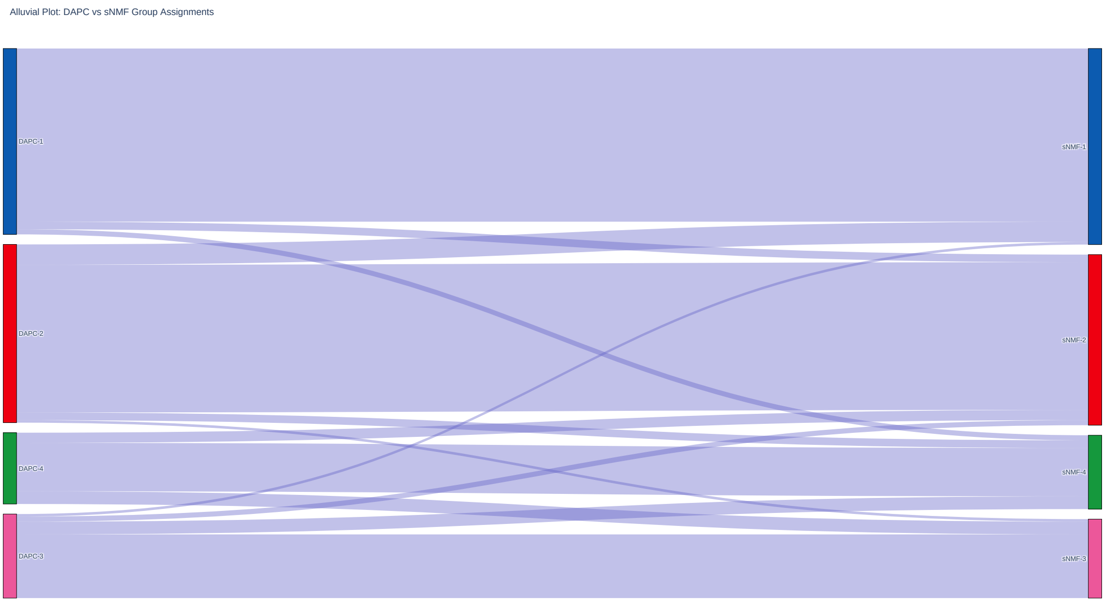


Figure S1. Concordance between sNMF and DAPC Groupings.
Alluvial (Sankey) diagram illustrating the correspondence of individual group assignments between sNMF and DAPC clustering approaches.
Flow widths represent the number of shared individuals.
Most individuals show consistent group membership across both methods.

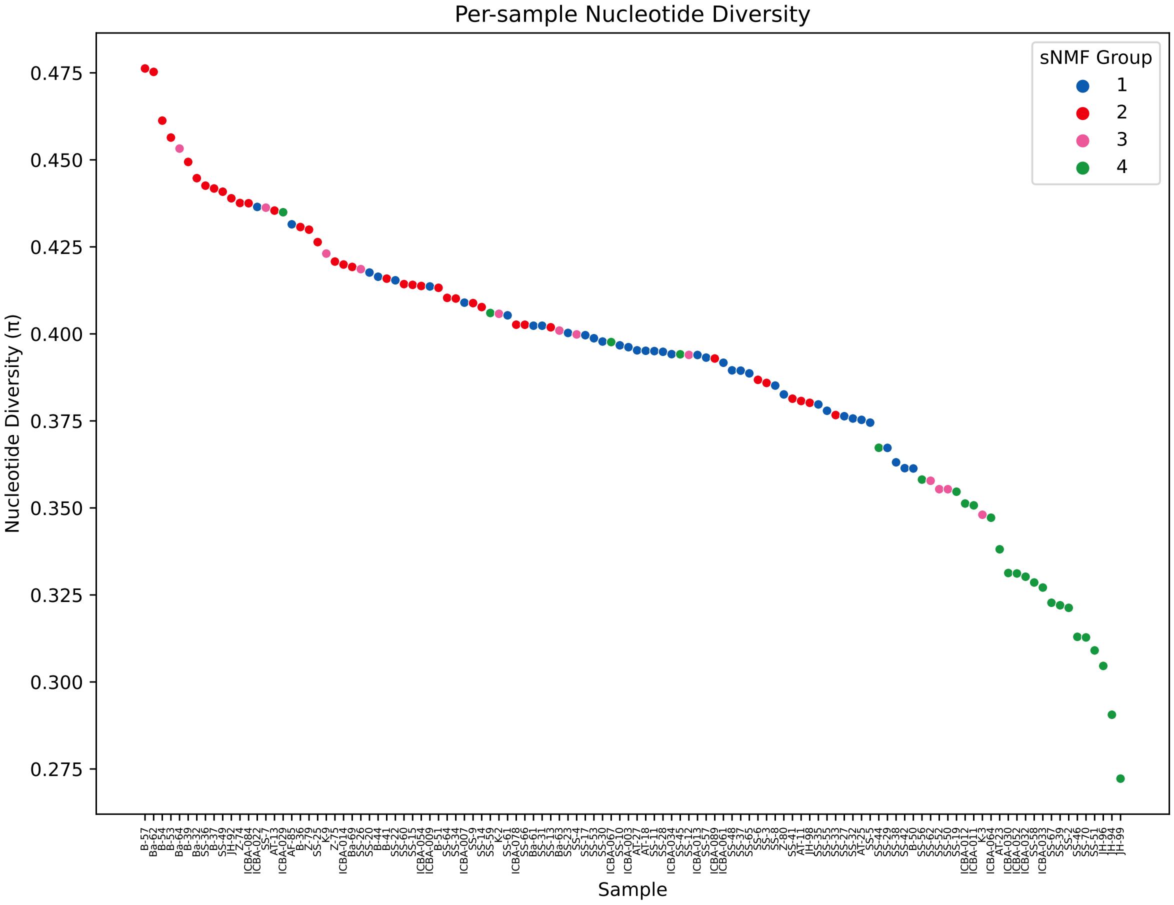


Figure S2. Per-Sample Nucleotide Diversity .
Violin plot depicting the distribution of per-individual nucleotide diversity , color-coded by sNMF group.
Highlights within-group and between-group genetic variation.

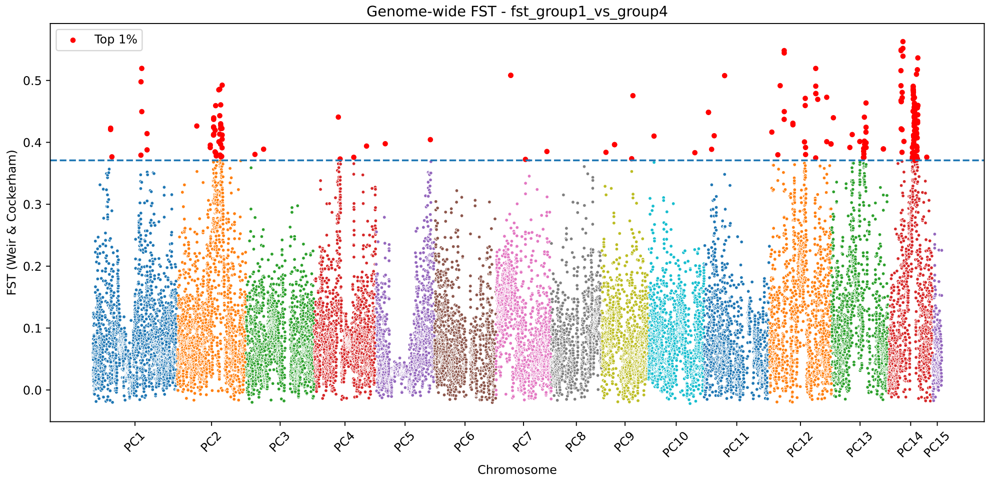


Figure S3. Genome-Wide $F_{ST}$ Scan: Group–1 vs Group–4.
Manhattan plot showing genome-wide $F_{ST}$ values in sliding windows.
The top 1% most differentiated regions are highlighted, indicating candidate loci for population divergence.

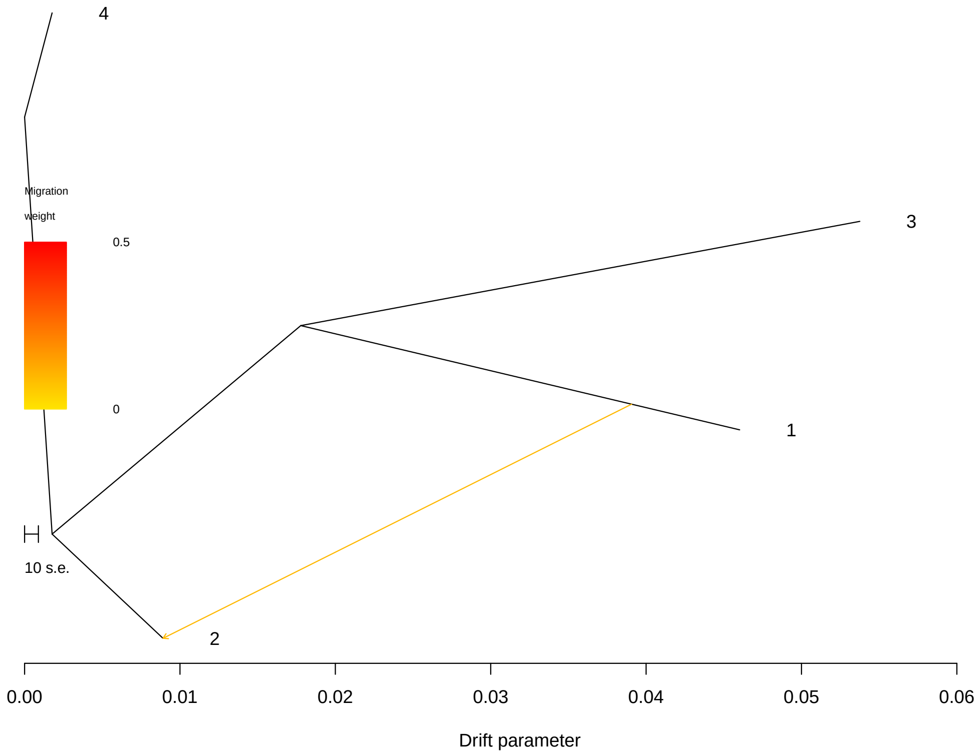


Figure S4. TreeMix Inference of Gene Flow Events.
Maximum likelihood population tree inferred using TreeMix, showing inferred migration edges among Ghaf genetic groups.
A notable migration event is detected from Group–1 to Group–2.

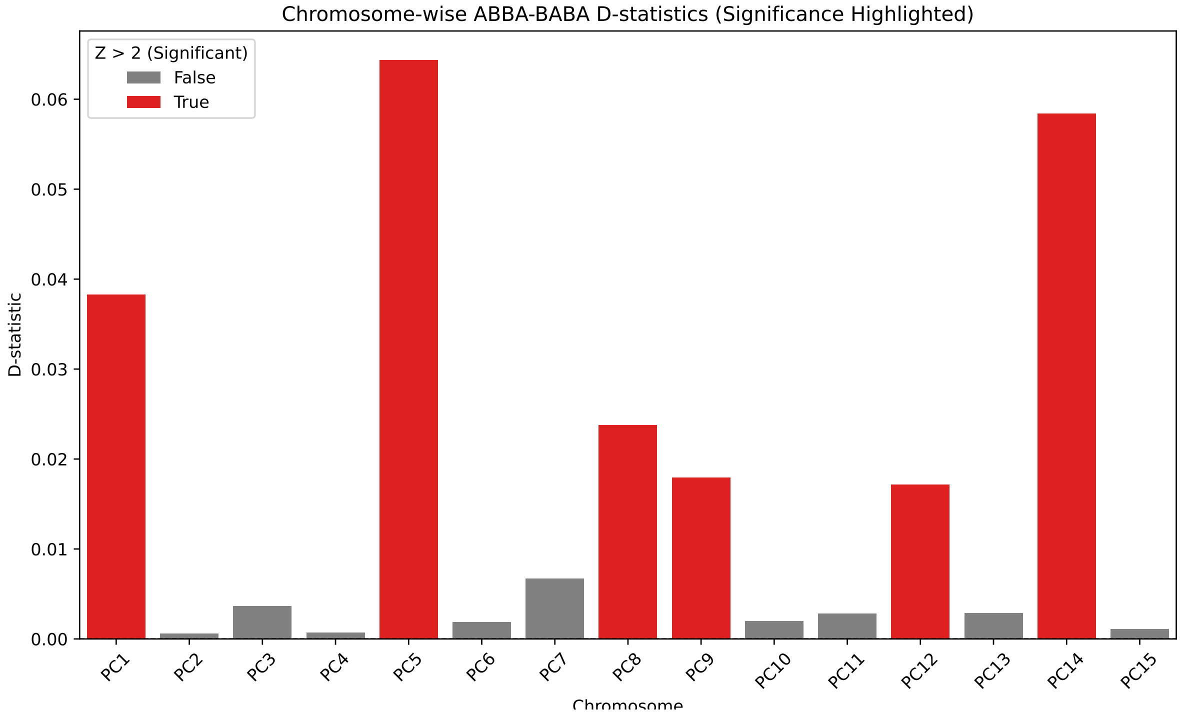


Figure S5. ABBA–BABA D–Statistic Test Across Chromosomes.
Barplot displaying chromosome–level D–statistics.
Significant introgression signals ($Z > 2$) are observed for chromosomes PC1, PC5, PC8, PC9, PC12, and PC14, indicating localized gene flow.

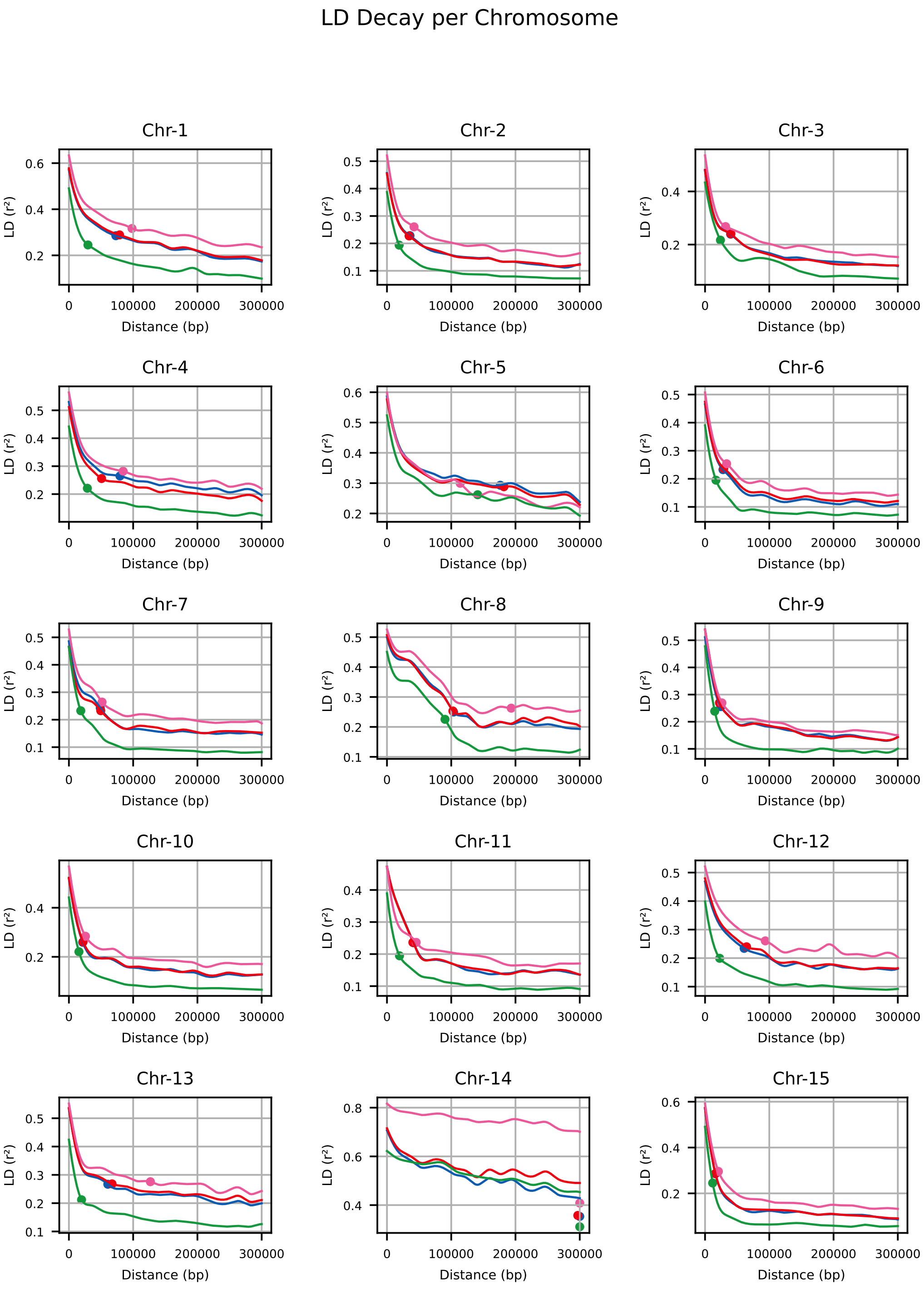


Figure S6. Linkage Disequilibrium (LD) Decay by Group and Chromosome.
Grid of LD decay curves ($r_2$ vs physical distance) for each sNMF group across all 15 chromosomes. Group–4 shows a more rapid decay pattern, consistent with elevated inbreeding and reduced diversity.
